# Clustering rare diseases within an ontology-enriched knowledge graph

**DOI:** 10.1101/2023.02.15.528673

**Authors:** Jaleal Sanjak, Qian Zhu, Ewy A. Mathé

## Abstract

**Objective:** Identifying sets of rare diseases with shared aspects of etiology and pathophysiology may enable drug repurposing and/or platform based therapeutic development. Toward that aim, we utilized an integrative knowledge graph-based approach to constructing clusters of rare diseases.

**Materials and Methods:** Data on 3,242 rare diseases were extracted from the National Center for Advancing Translational Science (NCATS) Genetic and Rare Diseases Information center (GARD) internal data resources. The rare disease data was enriched with additional biomedical data, including gene and phenotype ontologies, biological pathway data and small molecule-target activity data, to create a knowledge graph (KG). Node embeddings were used to convert nodes into vectors upon which k-means clustering was applied. We validated the disease clusters through semantic similarity and feature enrichment analysis.

**Results:** A node embedding model was trained on the ontology enriched rare disease KG and k-means clustering was applied to the embedding vectors resulting in 37 disease clusters with a mean size of 87 diseases. We validate the disease clusters quantitatively by looking at semantic similarity of clustered diseases, using the Orphanet Rare Disease Ontology. In addition, the clusters were analyzed for enrichment of associated genes, revealing that the enriched genes within clusters were shown to be highly related.

**Discussion:** We demonstrate that node embeddings are an effective method for clustering diseases within a heterogenous KG. Semantically similar diseases and relevant enriched genes have been uncovered within the clusters. Connections between disease clusters and approved or investigational drugs are enumerated for follow-up efforts.

**Conclusion:** Our study lays out a method for clustering rare diseases using the graph node embeddings. We develop an easy to maintain pipeline that can be updated when new data on rare diseases emerges. The embeddings themselves can be paired with other representation learning methods for other data types, such as drugs, to address other predictive modeling problems. Detailed subnetwork analysis and in-depth review of individual clusters may lead to translatable findings. Future work will focus on incorporation of additional data sources, with a particular focus on common disease data.

## Background and Significance

Rare diseases affect up to 25-30 million people in the US[1] and more than 300 million worldwide[2], making rare diseases common as a collective. The burden of rare disease is disproportionately high because patients living with rare diseases tend to incur high healthcare costs along the course of long diagnostic odysseys and intensive treatment regimens[3, 4]. Furthermore, the population of rare disease patients is distributed across 5,000-10,000 distinct diseases[5], yet the vast majority have no approved therapeutics. These factors create a clear and present need for research and development of new therapeutic options.

Methods that enable research and development efforts to make advances toward treatments for multiple diseases simultaneously may offer a path forward. Some such methods are already in practice, including therapeutic platforms like gene therapies[6], basket clinical trials[7] and drug repurposing[8]. Both basket clinical trials and drug repurposing require knowledge of the connections between diseases through their underlying causal factors.

Following similar efforts in the broader biomedical community[9], data integration and harmonization efforts in the rare disease space have emerged to support research and development aimed at multiple diseases at once. The Encyclopedia of Rare disease Annotations for Precision Medicine (eRAM) was built using a text-mining approach from the biomedical literature to connect and annotate diseases, genes, phenotypes into a system designed for use by clinicians[10]. The RDMap utilized multiple biomedical ontologies cluster diseases within a multidimensional map of rare diseases for researchers and clinicians to explore similarities amongst diseases[11]. Both eRAM and RDMap propose methods for calculating the similarity of rare diseases using phenotype and gene annotations individually and then combining the similarity scores.

The National Center for Advancing Translational Sciences (NCATS) supports the Genetic and Rare Diseases (GARD) Information Center to maintain data on rare diseases with the United States. A preliminary attempt was made to harmonize data across the GARD diseases using multi-source mappings across diseases and genes and phenotype annotations[12]. Here we follow up on that study and use the similarity between diseases, with respect to their position within our KG, to perform disease clustering. Three factors differentiate our study from prior efforts: 1) the incorporation of explicit biological pathway and small molecule activity data, 2) the focus specifically on diseases tracked by GARD, and 3) the use of graph node embeddings.

DeepWalk[13] and Node2Vec[14] are notable graph node embedding algorithms that convert graph nodes into vectors by applying word2vec[15] embedding models to random walks taken from across the knowledge graph (or any graph structured dataset). OPA2Vec[16] and subsequently DL2Vec[17] were developed for the specific application of graph node embedding methods to the biomedical domain, with a particular emphasis on the use of semantic ontologies. A recent study utilizes the structure of the gene ontology to create gene and disease embeddings using only gene interaction data and gene-disease annotations[18]. In this study, we derive graph node embeddings for disease nodes within a KG containing diseases directly connected to associated genes and phenotypes, and further enriched with small molecules (both drugs and metabolites), molecular pathway data, and biomedical ontologies. Our embeddings were generated using DL2Vec, modified to balance the probability of traversal from a disease-node to either a gene-node or a phenotype-node. By using a variant of DL2Vec, we implicitly capture the semantic information contained within the ontology structures alongside the direct connections between diseases and genes/phenotypes.

The disease-node embeddings are clustered, and the resulting clusters were analyzed for both validation and interpretation. We show that indeed graph node embeddings can be used to generate coherent, as measured through both quantitative and qualitative analysis, clusters of rare diseases within a heterogenous knowledge graph. Further, several of our rare disease clusters show promising connections to drugs and investigational compounds.

## Materials and Methods

### Data sources

Several data resources were integrated to construct a rare disease focused KG. The GARD internal data resources were queried to obtain the overlap between GARD and Orphanet disease lists. We focused on GARD diseases to support additional efforts sponsored by NCATS and relied on Orphanet as an external source of validity for the disease list. Gene and phenotype annotations for each disease were obtained from Orphanet’s ORPHADATA v4.0 resource[19]. The Gene Ontology[20, 21] (GO release 2021-10-26) and The Human Phenotype Ontology[22] (HPO release 2021-10-10) were both obtained from The OBO Foundry[23]. GO annotations for genes were obtained from NIH-NCBI (ftp://ftp.ncbi.nlm.nih.gov/gene/DATA/gene2go.gz). Metabolic, gene regulatory and physical interactions between genes, gene products, and small molecules were obtained from The Pathway Commons (PTC v12)[24]. Additional connections between genes and small molecules, based on published bioassay results, were obtained from Pharos v3.8.0[25, 26] via their API.

### Rare disease network construction

HPO and GO ontologies were extended to include additional logical connections with the ELK reasoner[27]. The HPO class of “HP:0000005: Mode of Inheritance” and its subclasses were pruned from the processed HPO ontology because the presence of this class created an overly connected network and does not reflect a relationship between diseases that is relevant to our use cases. The PTC and Pharos data were ingested as edge lists and harmonized through the ChEMBL-ChEBI and ChEBI-PubChem mapping files provided by UniChem(accessed on 2022-04-25)[28]. In cases where a particular entity mapped to multiple HPO or GO classes within a particular subtree of the ontology, e.g., a gene mapped to two GO terms that shared a parent-child relationship such as protein binding (GO:0005515) and kinase binding (GO:0019901), only the annotation to the lowest subclass, e.g., kinase binding, was kept. The pruning process removed redundant GO and HPO term annotations while preserving the annotations with the most semantic information content. Finally, all data were loaded into a single graph object.

### Graph node embeddings

Random walks emanating from each rare disease node were generated following a modified DL2Vec approach. The random walks were compiled into a corpus with each walk sequence. The length of each random walk and the number of random walks generated per disease were varied as part of our sensitivity analysis. Many diseases have far more HPO phenotype annotations than gene associations, yet the gene associations provide a very informative connection to the molecular processes involved in the disease. Therefore, to give more weight to the gene annotations, the probability of taking a random walk step was balanced between the gene and HPO annotations for diseases with both annotation types. Word2Vec was used to ingest the random walks and generate word embedding models for various combinations of walk length and walk count. We used the skip-gram Word2Vec architecture and varied the vector embedding dimension and the context window size.

Because we do not have a gold standard labelled dataset, we relied on internal clustering metrics to tune random walk and embedding model hyperparameters. We used three internal clustering metrics that each captures a different aspect of clustering quality: the silhouette score, which compares the intra-cluster distance to the nearest cluster distance for each sample; the davies-bouldin index, which compares the intra-cluster distance with the inter-cluster distances (lower is better); and the calinksi-harabasz index, which measures the within versus between cluster variances. Embedding models of different dimensions are not directly comparable with these internal clustering metrics. Therefore, we relied on heuristic rule as guidance coupled with empirical analysis of the complexity feature space captured in the embedding model. We started by assessing the fourth root of the total number of words in our corpus. In our case the number of words is the total number of unique nodes and unique edges traversed in the random walks, and results in a suggested embedding dimension of 26. Using this heuristic, we selected a range of embedding dimensions between 4 and 128 to test. We then performed sensitivity analysis with the other three parameters (the number of walks, the length of the walks and the embedding context window size) within each embedding dimension.

### Rare disease node clustering

The disease node embedding vectors were extracted from the Word2Vec models and concatenated to form a matrix where each row represents a disease, and each column is treated as a feature. We then applied K-means clustering using the python package scikit-learn[29] to cluster diseases. The number of clusters was selected using a variant of the elbow method entitled the kneedle algorithm[30] as implemented in the kneed v0.8.1 python package.

### Feature enrichment analysis

The node embedding process captures complex information regarding the local context surrounding a disease node within the knowledge graph, including phenotypes, genes, small molecules, (including drugs and metabolites) and more. This approach enables the clustering model to consider a rich feature space for each disease and is therefore more powerful than simply modeling direct annotations associated with each disease. However, this embedding process makes it more difficult to interpret in detail why a particular set of diseases ended up in a cluster together, yet this information is key for understanding the results and determine next steps. Therefore, we pursued two different forms of feature enrichment analysis.

First, we tested for gene enrichment for each cluster, with the aim of determining which genes were represented more frequently than expected by chance in each cluster. We counted the number of diseases associated with each gene within each disease cluster. To test against the null hypothesis that diseases were assigned to clusters independently of their gene annotations, the disease to cluster assignments were permuted 500,000 times, keeping the gene to disease annotations the same (noting that those are derived from the graph annotations and not our embeddings). The distribution of the counts of gene to cluster assignments for each gene within each cluster in the permuted data represent the null hypothesis of random grouping of genes within the clusters. A p-value was calculated based on the number permutations in which the counts of each gene within each cluster was greater than observed.

Random walk feature importance was estimated by calculating the term frequency-inverse document frequency (tf-idf)[31] of each feature within windows surrounding occurrence of each disease within the random walk corpus. The window size was selected to be consistent with the window size used in training the vector embedding model. The feature occurrences for each disease were summed within each cluster to obtain the term frequency. The presence or absence of each feature within each cluster was treated as the “document” frequency. An empirical cumulative distribution function was constructed from the tf-idf values and various percentile cutoffs were used to then summarize how informative each feature type for each cluster.

We sought to further interpret the known relationships amongst clustered diseases with respect to their enriched gene annotations. We therefore used STRINGDB v11.5[32] to 1) assess whether the enriched genes within each cluster have more connections had than expected by chance and 2) identification of enriched GO terms first for each set of genes.

### Semantic similarity validation

The Orphanet Rare Disease Ontology (ORDO v4.0) [33] was used to calculate pairwise semantic similarity amongst diseases based on the Sanchez information criteria[34]. We sampled 100 random sets of diseases of size 87 (the average number of diseases per cluster) to obtain a sampling distribution of average pairwise semantic similarity. We then performed a one sample Student’s t-test to statistically evaluate the difference in the mean average semantic similarity between the disease clusters and the random samples by assuming that the mean and variance parameters from the random samples represent the sampling distribution under the null hypothesis.

## Results

### Exploratory analysis of the rare disease network

The node degree (number of connections for each node) distribution of the knowledge graph is shown in Figure S1. The overall degree distribution of the knowledge graph is nearly linear on a log-log scale, indicative of a power-law distribution. The degree distribution is largely determined by the preponderance of small molecule nodes (N=348,395).

To evaluate the connectivity between diseases in the graph, we calculated the distribution of shortest path lengths between every pair of disease nodes and the results are shown in Figure S2. The most common path lengths were four and two corresponding to having three and one intermediate node, respectively. Genes and phenotypes are the only direct connections to diseases and both the genes and phenotypes are connected through the GO and HPO ontologies respectively. The hierarchical tree structure of the ontologies included in the network result in an increase in the frequency of paths of length 4 compared to paths of length 3 or 5. For example a common type of path between diseases goes as follows: disease, HPO phenotype, HPO class, HPO phenotype, disease; this type of path has an edge length of 4. The longest path between any two disease nodes in the graph was 6 and so we expect that disease nodes will occur frequently within the random walks.

### Optimization of embedding dimension

Using the optimized parameters, we performed principal component analysis (PCA) on the disease embedding vectors from models with different embedding dimensions. By plotting the explained variance as a function of the number of principal components, we can visually inspect the degree to which dimensionality of the embedding space can be reduced. We built models with embedding dimension of 4, 8, 16, 32, 64 and 128. We observed that for models with embedding dimension of 4, 8 or 16 there was no drop off in variance explained by successive PC’s indicating that these feature spaces could not be significantly reduced. In contrast, models with dimension of 32 or higher showed a large decrease in variance explained by higher PC’s suggesting that those feature spaces could be reduced (Figure S3). Therefore, an embedding dimension of 32 was selected for final analysis. It is useful to note that this embedding dimension is roughly consistent with that recommended by the fourth root heuristic.

### Disease clusters

A total of 3,242 diseases were used to construct a rare disease network that included data on genes, phenotypes, small molecules, biological pathways, and biomedical ontologies. The rare disease network contained 439,691 nodes and 2,716,895 edges (see the Supplemental Material for an exploratory data analysis of the network). Figure 1 illustrates workflow used create disease clusters from the rare disease network.

**Figure 1:**
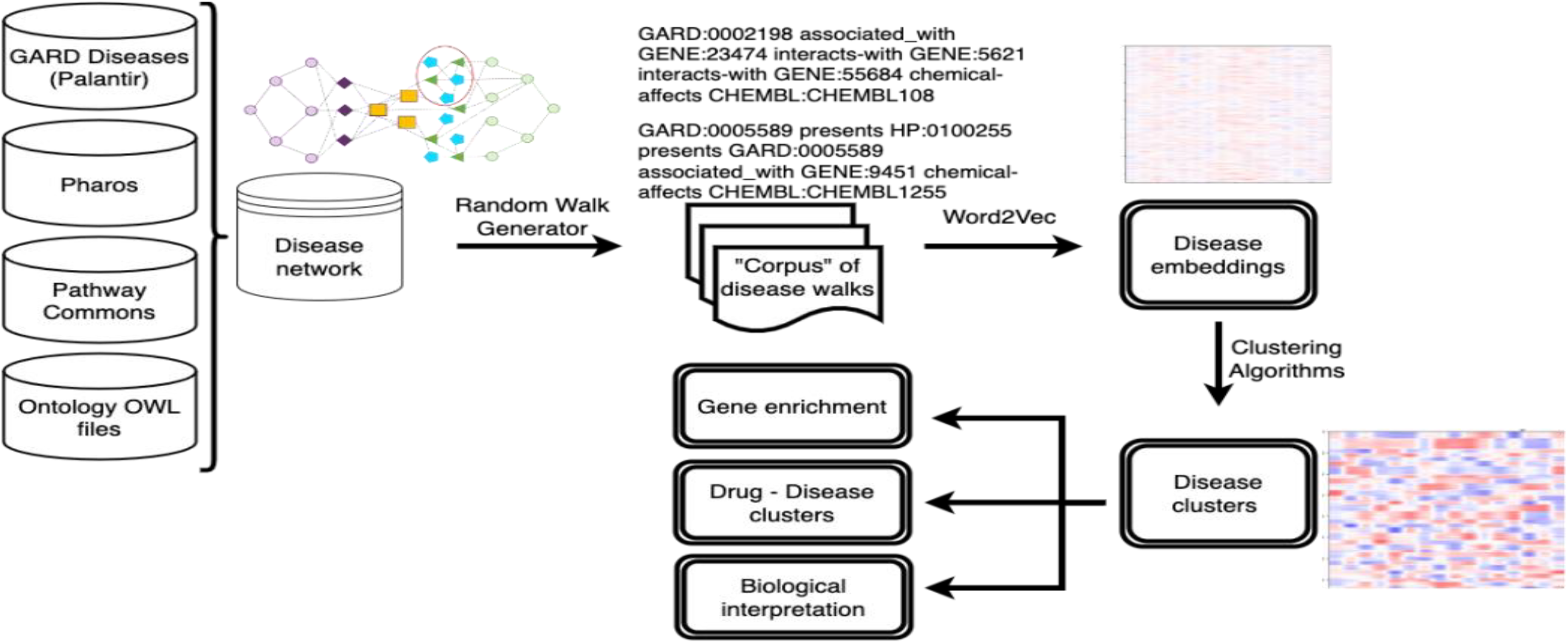
Rare disease clustering workflow. Input data resources are integrated into a single rare disease focused network. Random walks are performed to create a corpus of surveying the local context around each disease. Node embeddings are created and clustered. Post-hoc analyses are conducted to interpret and utilize the disease clusters

Several random walk and embedding model parameters were systematically varied to optimize the quality of the disease clusters produced. Parameters varied include the number of walks, the length of the walks, the embedding context window size, and the overall embedding dimension. Figure S4 shows several internal clustering metrics as a function of the number of walks per disease, across the various walk lengths and context window sizes for an embedding dimension of 32. The number of walks per disease had the largest effect on all clustering metrics. Performance increased with the number of walks with a plateau reached beyond 250 walks per disease. Based on this sensitivity analysis, we selected the following model parameters: 250 walks, walk length of 250 and context size of 20.

#### Final disease clustering model and its evaluation

Our final disease cluster model contained 37 clusters, with an average of 87 and a median of 83 diseases per cluster. The distribution of cluster sizes is shown in Figure S5. In Figure 2 the disease embedding values are plotted as a heatmap with diseases sorted either randomly or by cluster assignment, which shows that diseases with similar patterns across the embedding values tend to group together into clusters. To assess the degree of separation between the disease clusters we projected the disease clusters into a two-dimensional t-SNE map. The t-SNE projection of the disease clusters, colored by their cluster assignments, are depicted in Figure 3(a). The t-SNE map shows apparent separation amongst the disease clusters.

**Figure 2:**
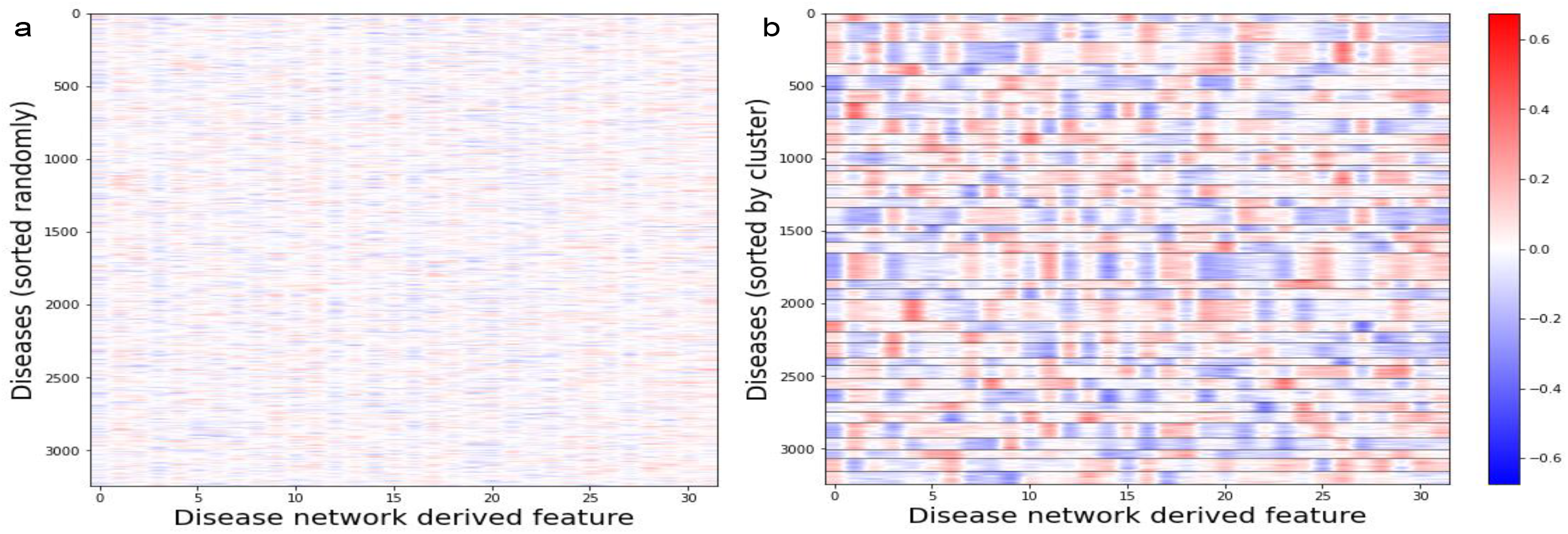
Disease embedding vectors are plotted in heatmaps (a)randomly sorted and (b)sorted by cluster. Each column in the heatmap corresponds to a dimension within the embedding vector space and each row corresponds to a disease. In figure b, the clusters are demarcated with horizontal black lines.

**Figure 3:**
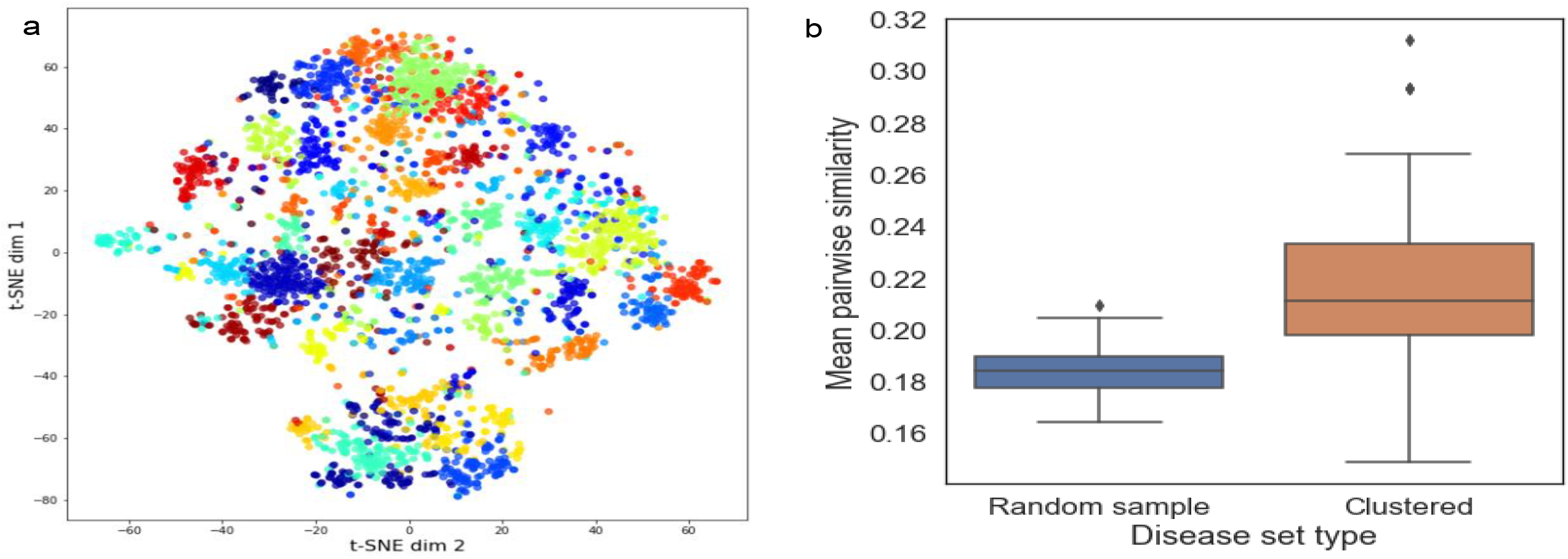
Visualizing and quantifying similarity within disease clusters. (a) t-SNE projections of the disease embedding vectors were created and plotted, points are colored according to their cluster membership. (b) The average pairwise Sanchez semantic similarity metric amongst disease within each cluster or within random samples.

To inspect the clusters quantitatively, we calculated a within-cluster semantic similarity index and compared that with randomly sampled disease sets. Specifically, we utilized the ORDO to calculate the Sanchez intrinsic information criterion within each disease cluster (see Methods). Figure 3(b) shows the distribution of the average semantic similarity both within each disease cluster and across a set of 100 randomly sampled diseases of size 87 (the average number of diseases per cluster). The t-test results suggest that the clustered diseases are significantly more semantically similar than a random selection of diseases (p=0.00024). These results indicate that the disease clustering model has captured information contained within the knowledge graph that is useful for assessing similarity amongst diseases.

### Cluster feature enrichment

Our first feature enrichment analysis focused strictly on direct genes annotations. By permuting the disease-cluster assignments we identified 585 genes enriched in at least one cluster at an FDR q-value cutoff of 0.01. We found that 12 genes were enriched within more than one cluster. The genes enriched in more than one cluster include genes associated with some major classes of diseases, such as: 1) oncogenes: KRAS, PTEN, TP53, KIT and FGFR1; 2) gene associated with musculoskeletal phenotypes: COL1A1, FKTN, GMPPB, POMT1, POMT2; 3) and genes associated with blood disorders: HBB, NPM1. The remaining 573 genes were enriched within only one cluster each (totaling 33 clusters with at least one enriched gene).

Second, we analyzed the random walks themselves to identify context features (e.g., genes, HPO terms, etc.) that make up each individual disease cluster. Figure 4 shows the number of nodes of each type as a function of the tf-idf percentile threshold for a set of clusters selected to be exemplary of different cluster archetypes. The cluster archetypes we identified include those almost exclusively dominated by HPO terms (clusters 16 and 19), those where the highest end of the td-idf distribution is dominated by genes (clusters 22 and 24), clusters with important GO terms and genes (cluster 7) and clusters with no features amongst the higher tf-idf percentiles (cluster 30).

**Figure 4:**
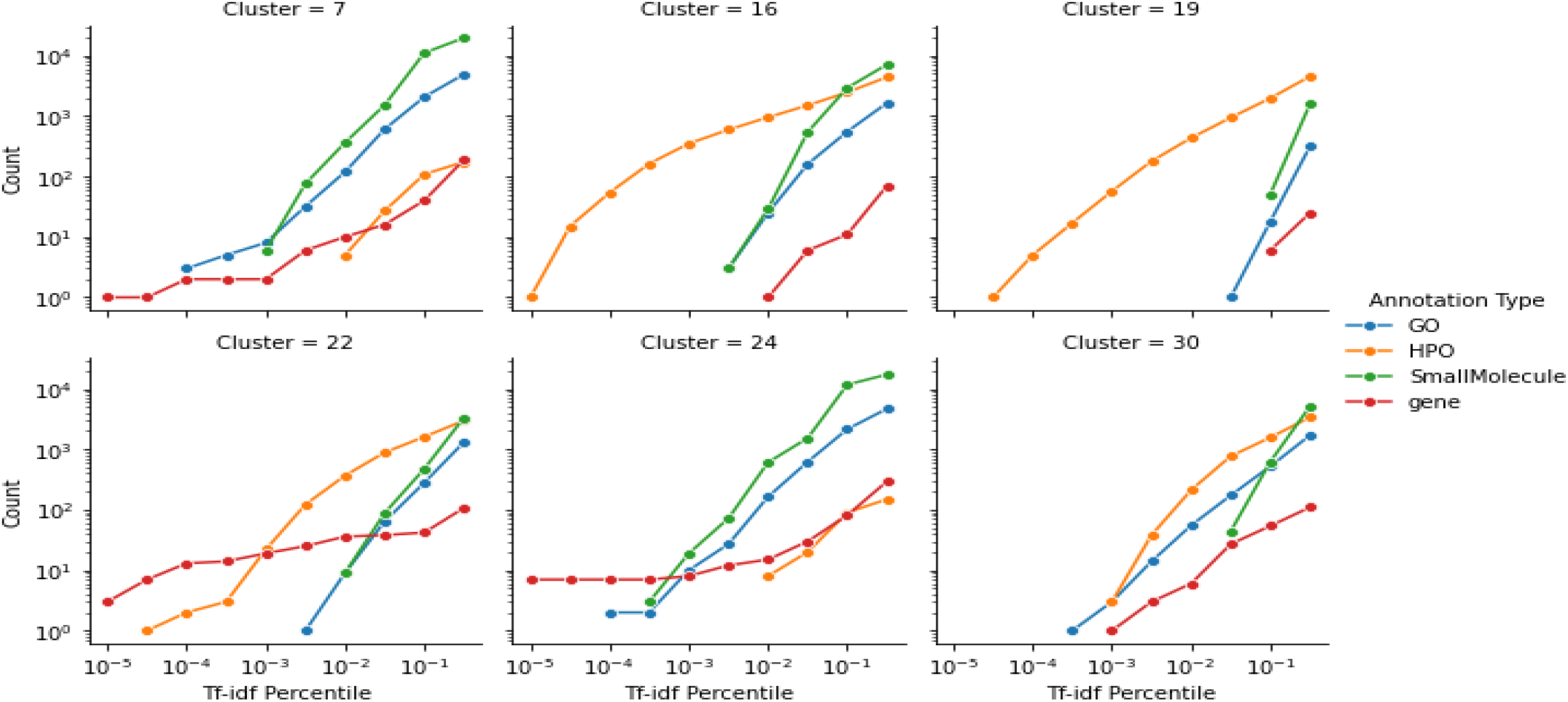
Counts by annotation feature type as a function of Tf-idf percentile threshold for a selected set of exemplary clusters.

### Interpreting the disease clusters

The sets of enriched gene annotations within each cluster were queried against STRINGDB. The enriched gene sets from 27 of the 33 clusters having at least one enriched gene had significantly more connections within STRINGDB than expected by chance. We manually reviewed the diseases, enriched genes, and enriched GO terms within each cluster with the goal of constructing concise descriptions of each cluster. Three clusters that could be concisely described are summarized in Table 1. Cluster 3 contains several neuromuscular and skeletal diseases such as Charcot-Marie Tooth disease type 2C and Duchenne Muscular Dystrophy. Cluster 7 is primarily composed of diseases that affect the visual system such as Cone-Rod Dystrophy and Leber Congenital Amaurosis. Cluster 28 contains several cardiac and electrophysiological diseases that are caused by mutations in ion channels (i.e., channelopathies). For all three clusters the enriched GO terms clearly align with the function of the associated genes and the etiology and pathophysiology of the diseases.

**Table 1.**
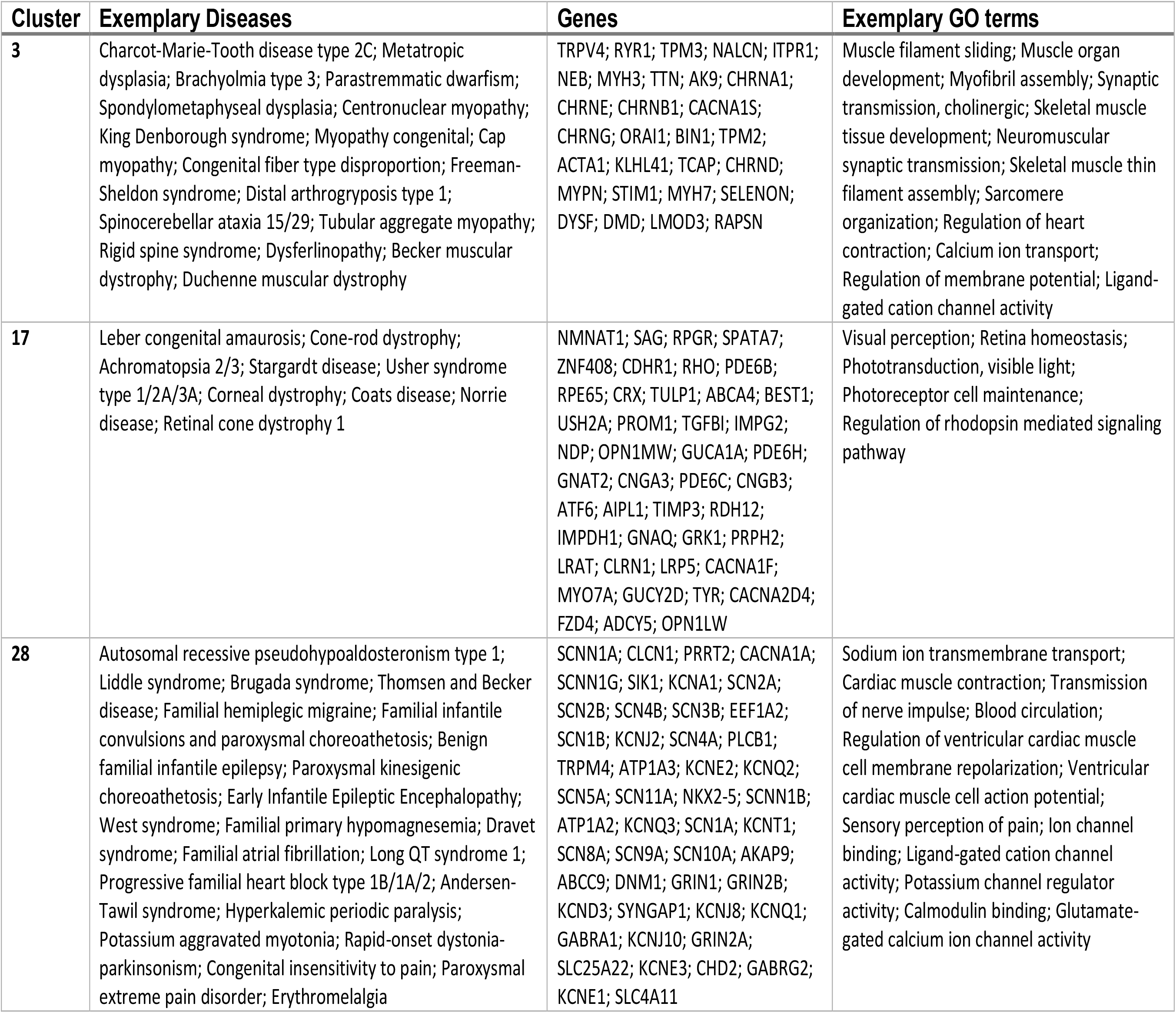
Example disease clusters with enriched genes and gene ontology terms

### Utility for drug repurposing

Drug repurposing is one important use case of our rare disease clusters. Identifying legitimate drug repurposing candidates will require in depth analysis of the clustered diseases and their connections to drugs. Here we sought to describe the connections between known drugs, gene targets, and the disease clusters.

Figure 5(a) shows the distribution of the number of gene targets by cluster and broken down by Target Druggability Level (TDL). The TDL is a qualitative label assigned to targets based on what is known about their chemistry, biology, and whether an approved drug is available[25, 26]. Tclin and Tchem rated targets are those with data indicative of druggability. Tclin rated targets have approved drugs with known mechanisms of action that target them and Tchem rated targets have ligands with activities known to be below target family specific thresholds. Tbio and Tdark rated targets are less obviously druggable; Tbio targets have published literature characterizing the target, yet no active ligands have been published and Tdark are considered understudied targets.

**Figure 5:**
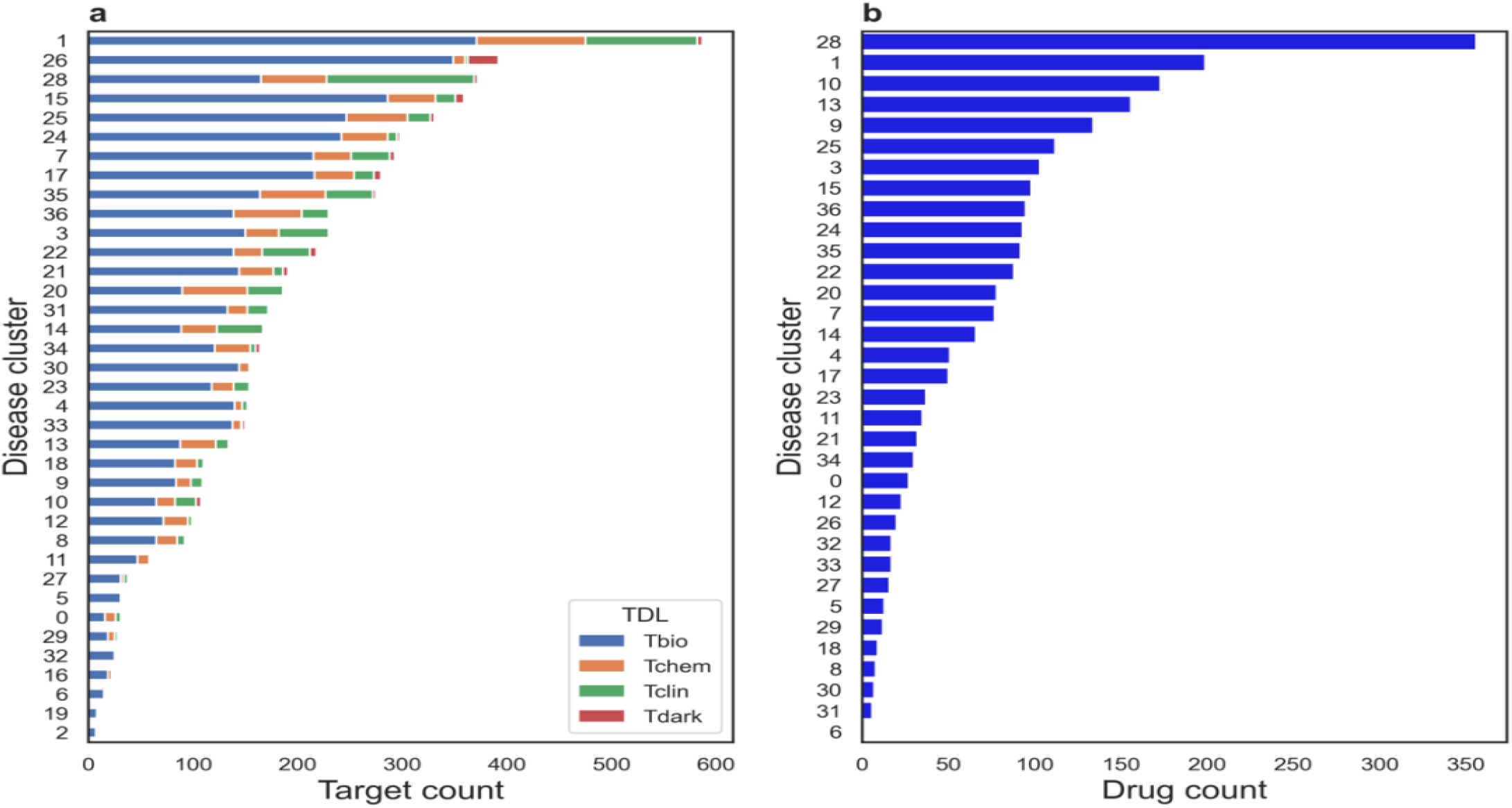
Summary of Pharos data by disease cluster. (a) The number of gene targets with a TDL label from Pharos within each cluster and (b) the number of drugs connected to each cluster through gene targets. Drugs were obtained from the Pharos based on their ‘Tclin’ ligand readiness level assignment.

At the time of writing this paper, the overall distribution of TDL values in Pharos was 11,867 Tbio, 5,932 Tdark, 1,930 Tchem and 685 Tclin. As in the overall TDL distribution, Figure 5(a) shows that Tbio is the largest category of gene targets in every cluster. In contrast to the overall TDL, relatively few targets with associations to the clustered rare diseases are rated as Tdark. The relative absence of Tdark targets is expected because a gene target will need to have literature support to be associated with one of the diseases in our network. Thirty-one out of 37 clusters have at least one gene target with a Tclin or Tchem rating, indicating that many of the clusters have putative drug repurposing candidates that could be mined in a more detailed analysis.

To explore the space of drug connections to our disease clusters, we filtered the original data from Pharos to include only approved drugs. Figure 5(b) shows the number of approved drugs with activity against a target in each cluster, based on the Pharos data. Among clusters with direct connections to drugs, the number of unique drugs per cluster ranged from 356 drugs in cluster 28 to just a single drug in cluster 6. In addition, clusters 2, 16 and 19 had no known drug connections. We anticipate using these summaries to help prioritize in depth follow-up studies on diseases within individual clusters.

## Discussion

We constructed a knowledge graph based on the overlap between rare diseases tracked by GARD and Orphanet. The graph is enriched with additional information on small molecules and biological pathways. We used this enriched network to construct graph node embedding vectors for each disease. Those embedding vectors were used as a feature matrix in a k-means clustering analysis. Hyperparameters of the embedding model and the k-means model were selected by a combination of heuristics, sensitivity analyses and explicit tuning. Our method identified 37 diseases clusters with an average of 87 diseases each. The quality of the resulting disease clustering model was validated by comparing semantic similarity within clusters compared to randomly selected disease sets, based on the ORDO, which was not part of our disease network. The semantic similarity of the clustered diseases combined with visual inspection of the cluster sorted embedding vectors and feature enrichment analysis suggests that our method has identified groups of diseases with features in common. Furthermore, manual review of the clustered diseases, enriched genes and enriched GO terms showed that many of the clusters are clearly composed of diseases related based on their causal gene and pathophysiology.

The use of semantic similarity within clusters as a form of validation begs the question: why was semantic similarity not the primary basis for clustering the diseases in the first place? Our aim was to expand beyond semantic ontological organization of diseases by a) directly using data related to the diseases and b) using higher order relationships between the diseases across data modalities. The former justification was taken up by both eRAM [10] and the RDMap [11] projects, which developed similarity scores using both Phenotype-HPO and Gene-GO linkages to rare diseases. Our approach expands upon those methods by integrating the Phenotype-HPO, Gene-GO and additional datasets into a single network. We capture higher order relationships amongst diseases by building graph node embeddings. The graph node embeddings provided and integrated representation of the network context of each disease across the heterogenous data types present in the network. However, our results are only as comprehensive as the underlying input data. Various sources of bias, including publication bias toward more prevalent diseases, limit the generalizability of our results. As more data is shared in the rare disease research space, our ability to extract insights from integrative data resources will increase.

Another key limitation of our study is the absence of common diseases. Most of the biomedical data pertains to common diseases. Therefore, expanding our knowledge graph to incorporate common disease information would greatly increase the scope and translational relevance of the work. However, the expansion of the graph would also create some challenges surrounding data source selection and overwhelming of the rare disease signal. Nevertheless, one goal for future work will be the incorporation of common disease data into the analysis.

Our analysis creates an additional layer of structure onto the large pool of rare diseases. We hope this structure will help strengthen drug repurposing efforts by enabling focus onto smaller disease sets. Yet it must be recognized that our analysis on its own does not directly yield translatable results; this is in part due to the limited interpretability of the graph node embeddings. In depth follow up, such as detailed subnetwork analysis or literature review will be required to take full advantage of our work – a task which will be taken up in a related manuscript.

## Conclusion

Our approach expands upon prior efforts to identify similarity of rare disease by integrating multiple data types and considering the higher order structure of the rare disease network simultaneously. We show that diseases in the clusters are enriched for similar gene annotations and that there are many possible connections to approved and investigational drugs. Future work will focus on expanding the knowledge graph with common disease data and detailed subnetwork analysis of the most promising clusters. At a higher level, we hope that our work shows the benefit of continued to growth of data sharing and integration within the rare disease research community.

## Acknowledgements

This research was supported in part by the Intramural Research Program of the National Center for Advancing Translational Sciences (NCATS), NIH under project ZIC TR000410-03. I would like to thank Dac-Trung Nguyen, Yanji Xu and Andy Patt for their insights during the formation of this project.

## Code and Data Availability

All code for executing analysis and constructing figures is contained here: https://github.com/jsanjak/RD-Clust. Publicly available data used in our workflow are referenced in scripts within the GitHub repository. GARD data used are presently not accessible to the public and are therefore provided as CSV files within the GitHub repository.

## Supplemental Figures

**Figure S1.**
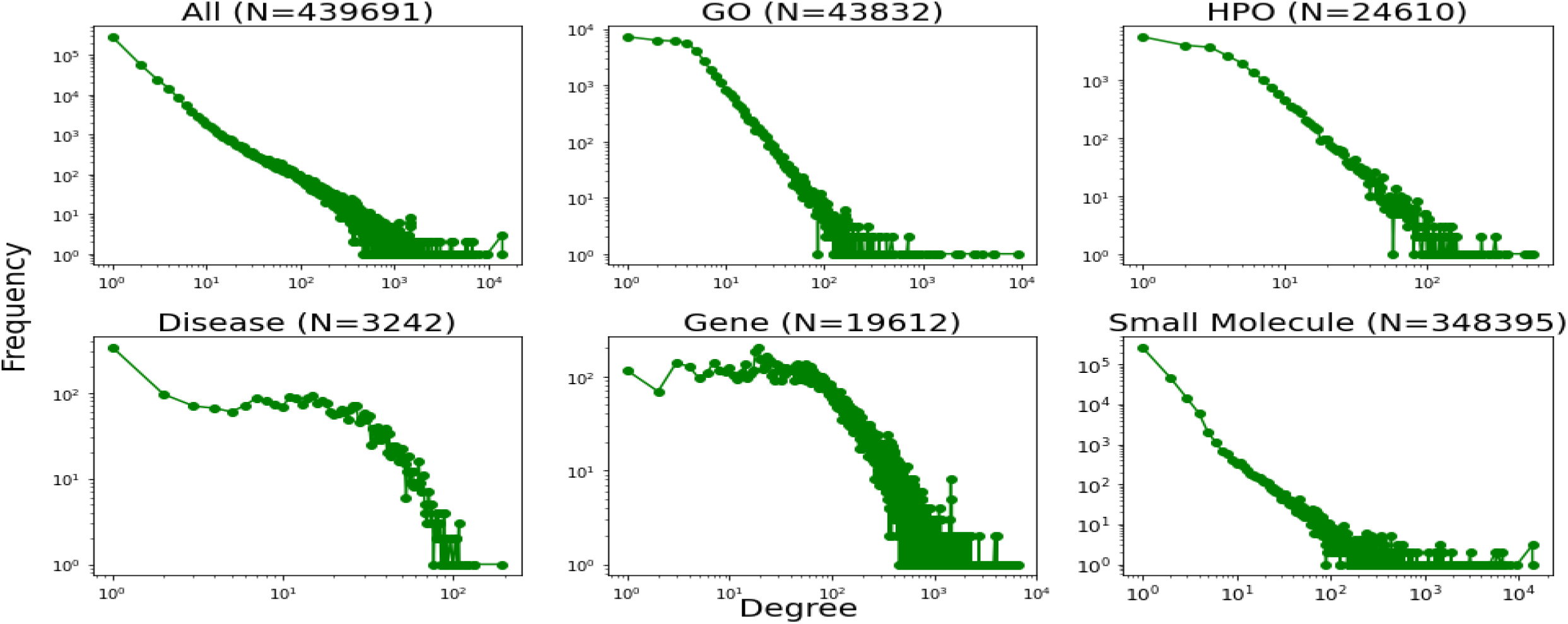
The degree distribution of various node types within the rare disease knowledge graph. Both y and x axes show the raw data values on a log-scale.

**Figure S2.**
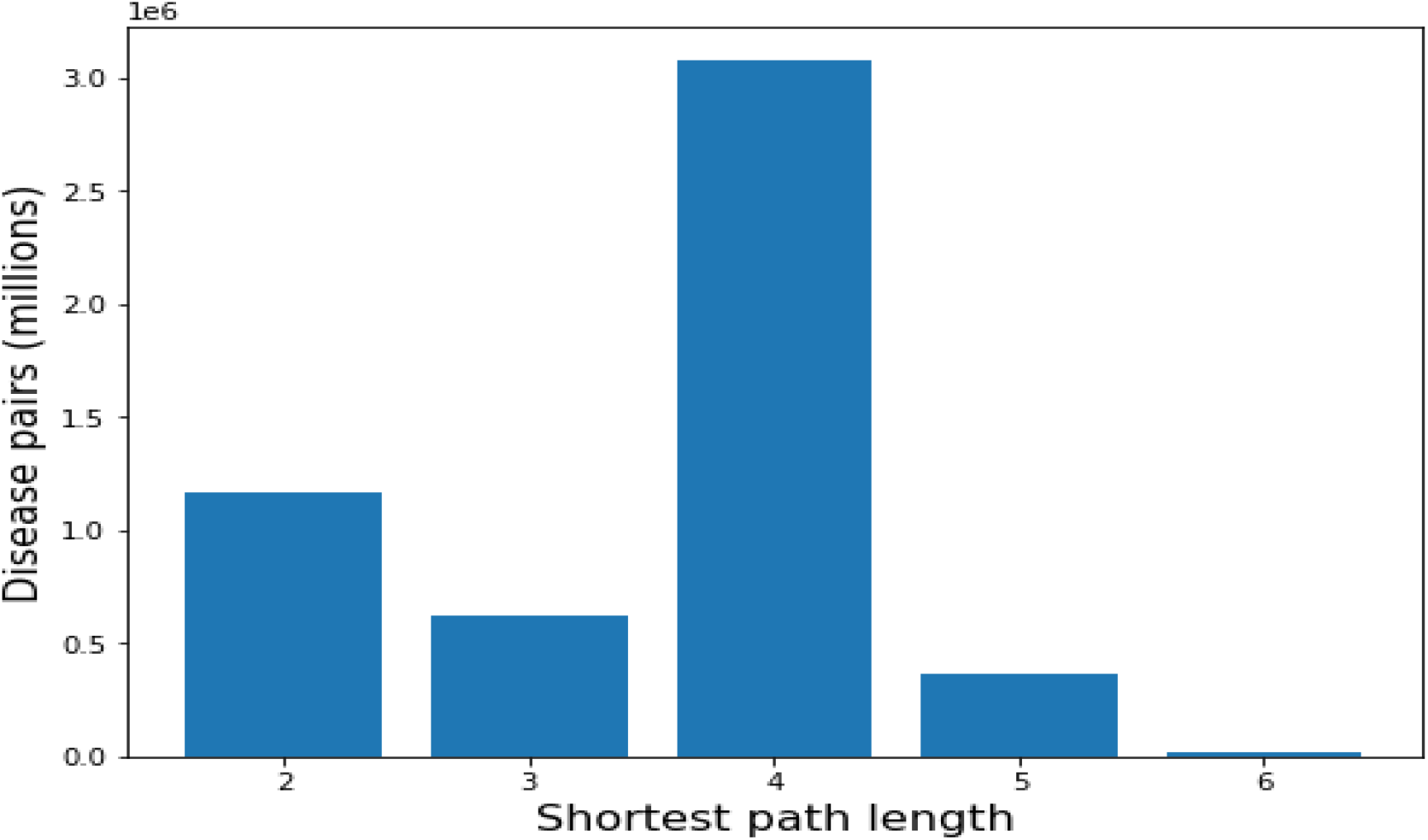
The distribution of shorest path lengths between all disease pairs within the rare disease knowledge graph

**Figure S3.**
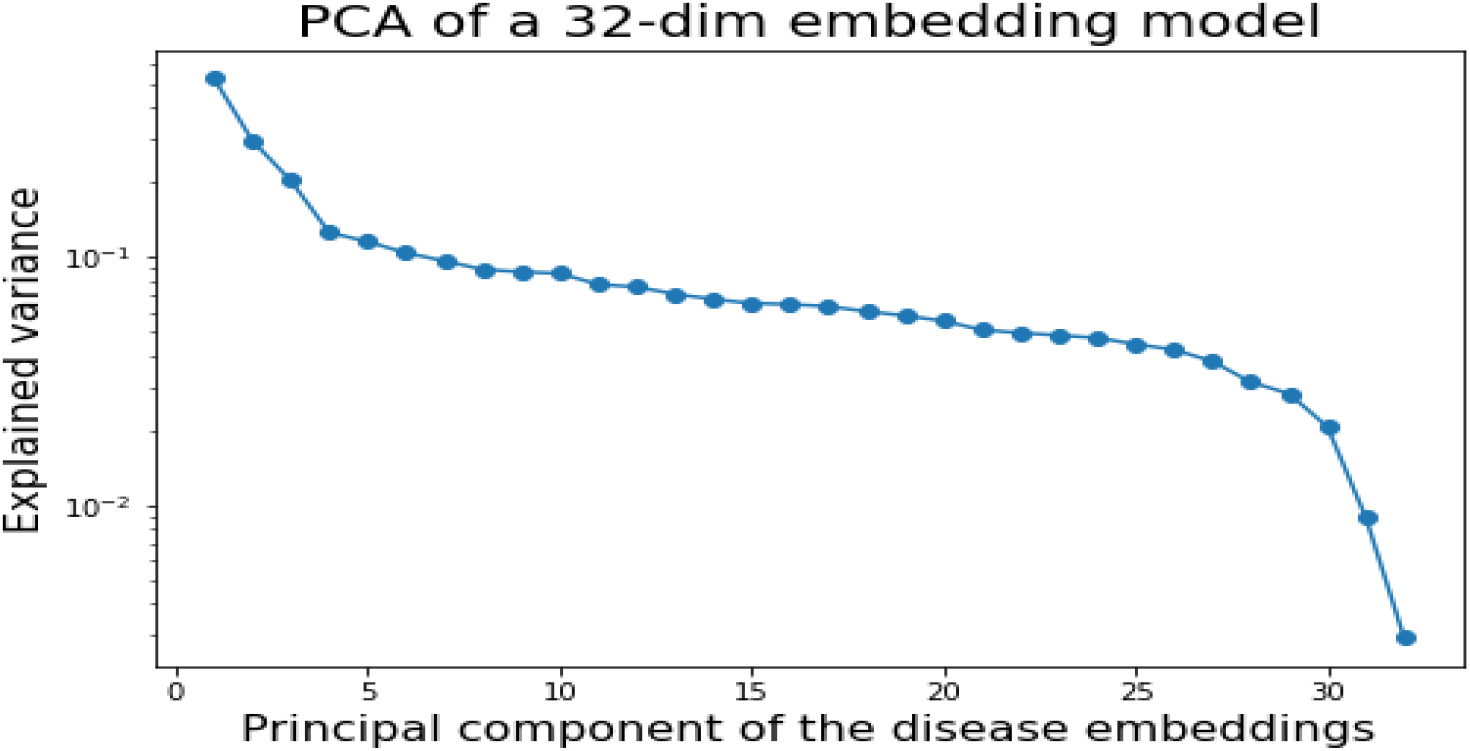
The variance in embedding values explained by successive principal component of the disease embedding matrix for an embedding dimension of 32.

**Figure S4:**
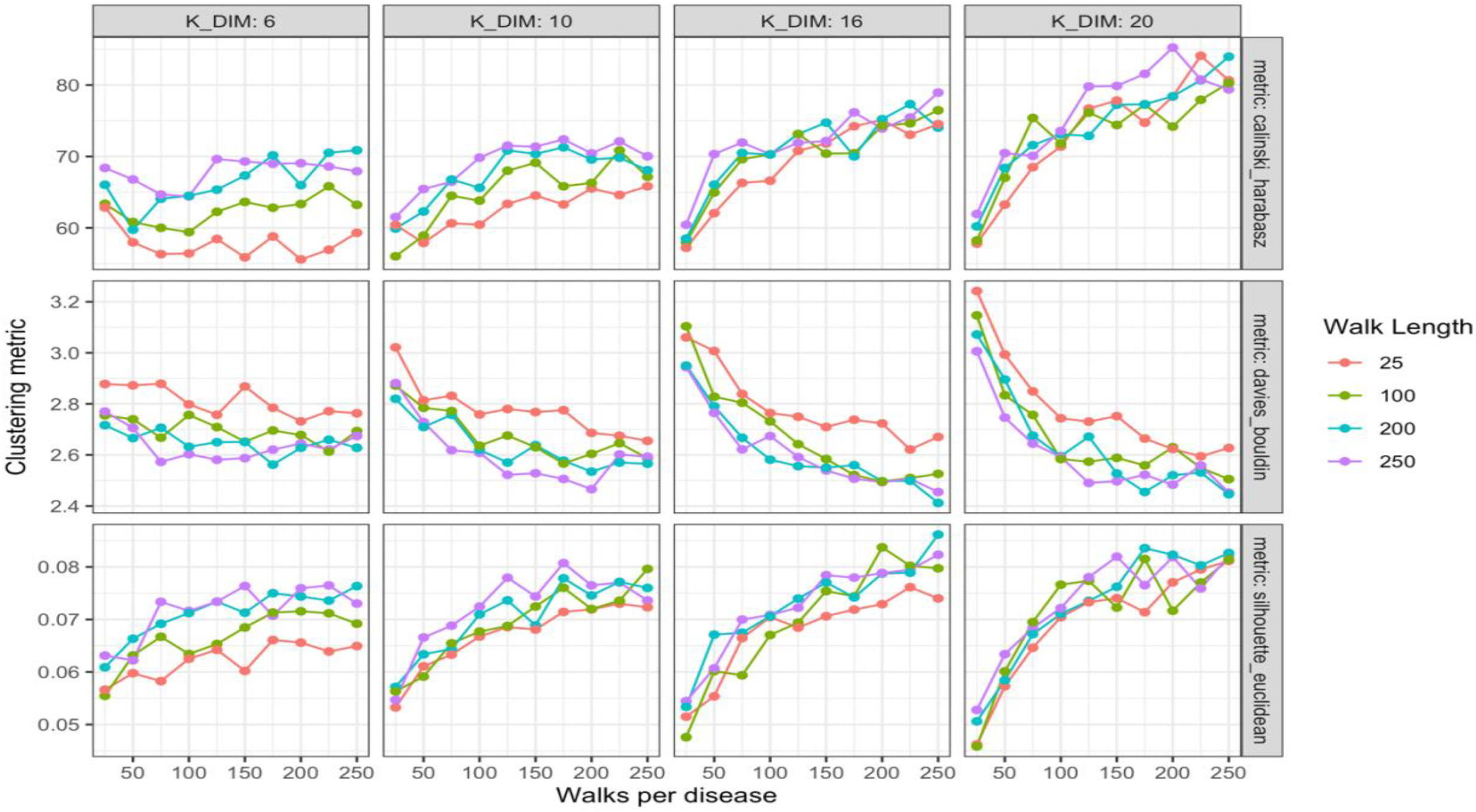
Internal clustering metrics as function of node embedding model hyper parameters. The colors of each line signify the length of the random walks used to construct the disease corpus. The chart columns vary the embedding context window (K_DIM). The chart rows show the three different clustering metric used, which are each plotted over the number of random walks generated per disease.

**Figure S5.**
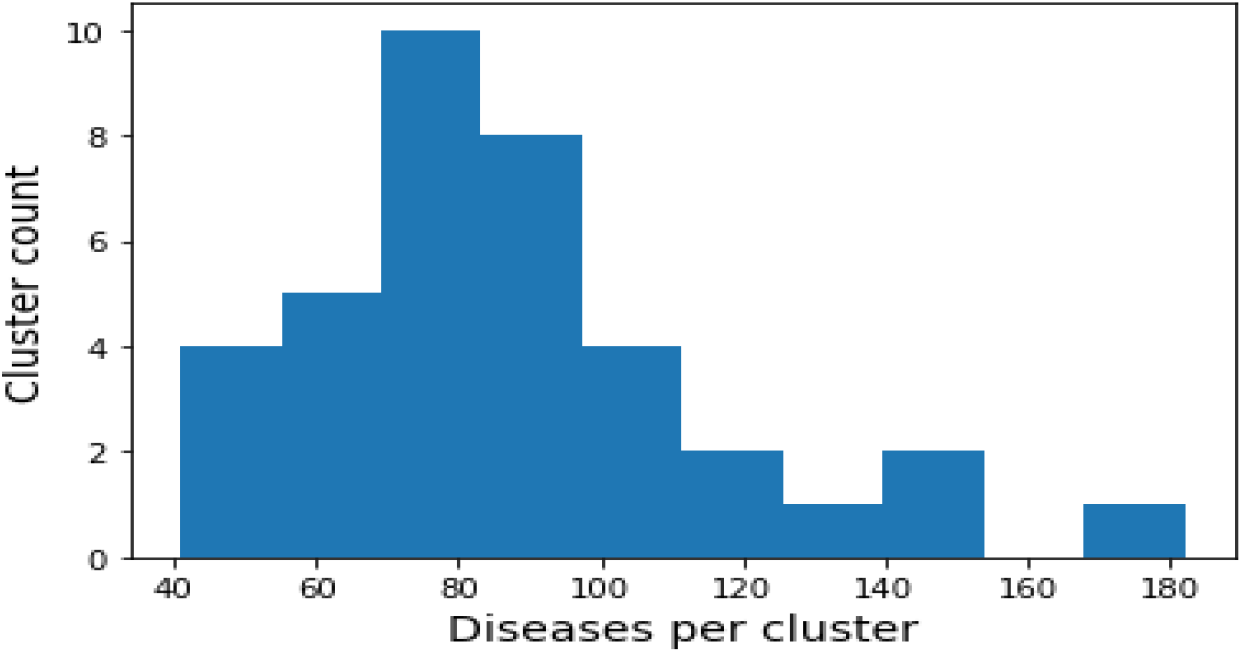
A histogram displaying the distribution of the number of diseases in each of the final clusters

## Notes

https://github.com/jsanjak/RD-Clust

### Competing Interest Statement

The authors have declared no competing interest.

